# Harnessing Interpretable Deep Learning to Predict Resistance in *Klebsiella pneumoniae*

**DOI:** 10.1101/2025.11.04.686393

**Authors:** Nicolas da Matta Freire Araujo, Márcia da Silva Chagas, Mateus Fernandes Santos, Renata Freire Alves Pereira, Rafaela Correia Brum, Felipe Ramos Pinheiro, Melise Chaves Silveira, Felicita Mabel Duré, Beatriz de Lima Alessio Müller, Audrien Alves Andrade de Souza, Alessandra Beatriz Santos Rondon Souza, Ágatha Ferreira de Souza, Ana Paula D’Alincourt Carvalho-Assef, Aline dos Santos Moreira, Marcelo Trindade dos Santos, Adriano Maurício de Almeida Côrtes, Bruno de Araújo Penna, Thiago Pavoni Gomes Chagas, Fábio Aguiar-Alves, Fabrício Alves Barbosa da Silva

**Affiliations:** Scientific Computing Program, Oswaldo Cruz Foundation (FIOCRUZ), Rio de Janeiro, Brazil; Department of Applied Mathematics, Institute of Mathematics, Federal University of Rio de Janeiro (UFRJ), Rio de Janeiro, 21941-909, Brazil; Laboratory of Molecular Epidemiology and Biotechnology, School of Pharmacy/Fluminense Federal University, Mario Viana, 523 - Niteroi/RJ, 2424.241-000, Brazil; Graduate Program in Microbiology and Parasitology. Biomedical Institute - Fluminense Federal University, Valonguinho Campus, Centro, Niteroi/RJ, 24020-140, Brazil; Department of System Engineering and Computation, State University of Rio de Janeiro, Rio de Janeiro, Brazil; Graduate Program in Pathology, Fluminense Federal University, Niteroi /RJ, Brazil; Bioinformatics Laboratory, National Laboratory for Scientific Computing (LNCC), Petrópolis, Brazil; Central Public Health Laboratory (LCSP), Ministry of Public Health and Social Welfare MSPyBS, Asunción, Paraguay; Laboratory of Applied Genomics and Bioinnovations - IOC/FIOCRUZ, Next-Generation Sequencing Platforms IOC/RPT01J - Network of Technological Platforms/FIOCRUZ, Brazil; Laboratory of Gram-Positive Cocci / Fluminense Federal University, Niteroi/RJ, Brazil; Laboratory of Bacteriology Applied to One Health and Antimicrobial Resistance, Oswaldo Cruz Foundation (FIOCRUZ), Rio de Janeiro, Brazil; Department of Computational Modeling, National Laboratory for Scientific Computing (LNCC), Petrópolis, Brazil; Systems Engineering and Computer Science Program, Coordination of Postgraduate Programs in Engineering (COPPE), Federal University of Rio de Janeiro (UFRJ), Rio de Janeiro, 21941-972, Brazil; Department of Pathology, School of Medicine, Fluminense Federal University, Niteroi/RJ, Brazil; Department of Pharmaceutical Sciences, Lloyd L. Gregory School of Pharmacy, Palm Beach Atlantic University – USA

**Keywords:** Deep learning, Genomics, *Klebsiella pneumoniae*, Modern Hopfield Networks

## Abstract

Antimicrobial resistance constitutes an escalating global health threat, complicating therapeutic management and increasing morbidity and mortality. Deep learning approaches have emerged as promising tools for bacterial profiling based on omics data, particularly for predicting antimicrobial susceptibility from genomic information. This task relies on identifying genomic signatures associated with resistance mechanisms. Here, DeepMDC is introduced as a deep learning architecture designed for bacterial profiling using whole-genome data. Given that precise annotation at the gene or mutation level is often costly and ambiguous, phenotypic classification is formulated as a Multiple Instance Learning (MIL) problem, in which each genome is represented as a bag of instances with a single associated label. The core of DeepMDC is a Modern Hopfield Network that processes all open reading frames (ORFs), including small ones, derived from genomic data. A key feature of the architecture is its interpretability, enabled by attention mechanisms that facilitate biological insight and hypothesis generation. The model was evaluated against *Klebsiella pneumoniae* and four clinically relevant antibiotics (meropenem, cefepime, ceftazidime, and gentamicin), achieving strong performance in several metrics. Notably, genes associated with resistance consistently received high attention scores during inference, which validates the architecture and eventually may generate new hypotheses.

## INTRODUCTION

Antimicrobial resistance represents a major global health challenge, increasing treatment complexity and contributing to higher morbidity and mortality rates. Deep learning approaches have shown strong potential in addressing this issue, particularly in bacterial profiling using omics data (Ardila et al., 2024). One key application is predicting antimicrobial susceptibility from whole-genome sequencing.

The classification of bacterial isolates as resistant or susceptible is based on phenotypic susceptibility testing, while the identification of genomic determinants reflects the potential for resistance rather than a definitive resistant phenotype. Conventional bioinformatics approaches rely on curated databases and rule-based tools such as PointFinder and ResFinder, which link specific mutations and acquired genes to resistance phenotypes (Aytan-Aktug et al., 2020). More recently, machine learning methods, including logistic regression, neural networks, and random forests, have demonstrated improved predictive performance (Aytan-Aktug et al., 2020; Peng et al., 2022).

Bacterial genomes exhibit high complexity and dimensionality, encompassing conserved core regions and highly variable accessory elements that contribute to resistance. Relevant genomic features include single-nucleotide polymorphisms, insertions and deletions, gene presence/absence patterns, and structural variations. These characteristics require computational models capable of handling heterogeneous and sparse data.

Advances in high-throughput sequencing technologies have significantly expanded the availability of genomic data, underscoring the need for methods to extract biologically relevant signals from large-scale datasets with limited supervision. In genomics, detailed annotation at the nucleotide or gene level is often unavailable or uncertain, and labels are typically assigned at the genome level (e.g., phenotype or taxonomy). Consequently, the contribution of individual genomic elements remains largely unannotated.

Multiple Instance Learning (MIL) provides a suitable framework for such scenarios. In MIL, data are organized into bags of instances, with labels assigned only at the bag level. This paradigm has been successfully applied in areas such as computational pathology, metagenomics, and immune repertoire analysis (Kraus et al., 2016; Hou et al., 2016; Rahman et al., 2020; Widrich et al., 2020). In genome-based classification, each genome can be modeled as a bag of instances (e.g., ORFs, k-mers), enabling the identification of discriminative patterns without requiring instance-level annotations.

While MIL provides a natural framework for weakly supervised genomic classification, most existing approaches remain focused on predictive performance, often treating genomic elements as independent contributors. However antimicrobial resistance is increasingly understood as an emergent property arising from interaction of multiple genomic components, including mobile genetic elements, regulatory regions, and structural variations (Ferrand et al., 2020). Therefore, models that capture not only the presence of features but also their relative importance within a genomic context are particularly valuable.

Recent advances in deep learning have introduced attention-based mechanisms that allow models to highlight informative regions of the input (Vaswani et al., 2017; Ramsauer et al., 2020). Nevertheless, their biological interpretation remains challenging, especially in genomic settings where annotation is incomplete or uncertain (Koo & Eddy, 2019; Yue et al., 2023). This gap motivates the development of approaches that combine weak supervision, interpretability, and the ability to capture distributed genomic signals.

Carbapenemase-producing *K. pneumoniae* has been classified by the World Health Organization as a critical priority pathogen (WHO, 2024). As a member of the ESKAPE group, *K. pneumoniae* is associated with severe clinical outcomes, including increased mortality, prolonged hospitalization, and elevated healthcare costs (De Oliveira et al., 2020; Loyola-Cruz et al., 2023). This opportunistic pathogen causes a range of infections, including urinary tract infections, pneumonia, bacteremia, and liver abscesses (Wang et al., 2020). Its resistance profile includes the production of β-lactamases and intrinsic resistance mediated by the *bla*SHV gene (Holt et al., 2015; Wyres et al., 2020). Virulence factors such as lipopolysaccharides, capsular polysaccharides, siderophores, and fimbriae further contribute to pathogenicity. Based on these characteristics, *K. pneumoniae* was selected as a model organism to evaluate DeepMDC, alongside four clinically relevant antimicrobial agents. Cefepime and ceftazidime are broad-spectrum β-lactams (fourth- and third-generation cephalosporins, respectively) with strong activity against Gram-negative bacteria, although their efficacy may be compromised by the production of extended-spectrum β-lactamases (ESBLs). Gentamicin, an aminoglycoside, exhibits rapid bactericidal activity by inhibiting protein synthesis. Meropenem, a carbapenem, is one of the most potent options for severe infections, particularly those caused by ESBL-producing strains, due to its stability against most β-lactamases and broad-spectrum activity. Together, these antibiotics are critically important for the clinical management of *K. pneumoniae*, as they encompass diverse modes of action and resistance mechanisms. DeepMDC achieved performance above 0.9 in several evaluation metrics. Furthermore, attention-based analysis revealed biologically meaningful patterns associated with antimicrobial resistance, supporting both predictive accuracy and interpretability.

In this work, we propose DeepMDC, a deep learning architecture for bacterial profiling from whole-genome data that frames the problem as an MIL problem. Specifically, given a genome, the model predicts if the strain is resistant or susceptible to a predefined antimicrobial agent. A key feature of DeepMDC is interpretability. During inference, DeepMDC assigns attention weights to each ORF in a genome, highlighting the ones most relevant to classification. Despite the correlational nature of attention coefficients, their interpretation is crucial for validating the model and eventually generating new insights into bacterial resistance.

## MATERIALS AND METHODS

### TRAINING DATA SELECTION AND PRE-PROCESSING

We recovered biosample accessions and associated resistance/susceptibility profiles for meropenem (MEM), gentamicin (GEN), ceftazidime (CAZ), and cefepime (FEP) from clinical *K. pneumoniae* strains described by Lueftinger et al. (2021). Because phenotypic annotations were not available for all antimicrobial agents across all isolates, antimicrobial-specific datasets were constructed, yielding distinct sample totals: 1,705 isolates for MEM, 2,145 for GEN, 1,975 for CAZ, and 1,908 for FEP. For each dataset, samples were partitioned into mutually exclusive training and testing sets using a 10:1 ratio. This approach ensured that the testing samples remained entirely independent during the model development phase. The distribution of samples between training and test sets for all antibiotics tested within this study is in Table 1. Sequencing reads for all selected samples were retrieved from the NCBI Sequence Read Archive (SRA) database.

**Table 1.**
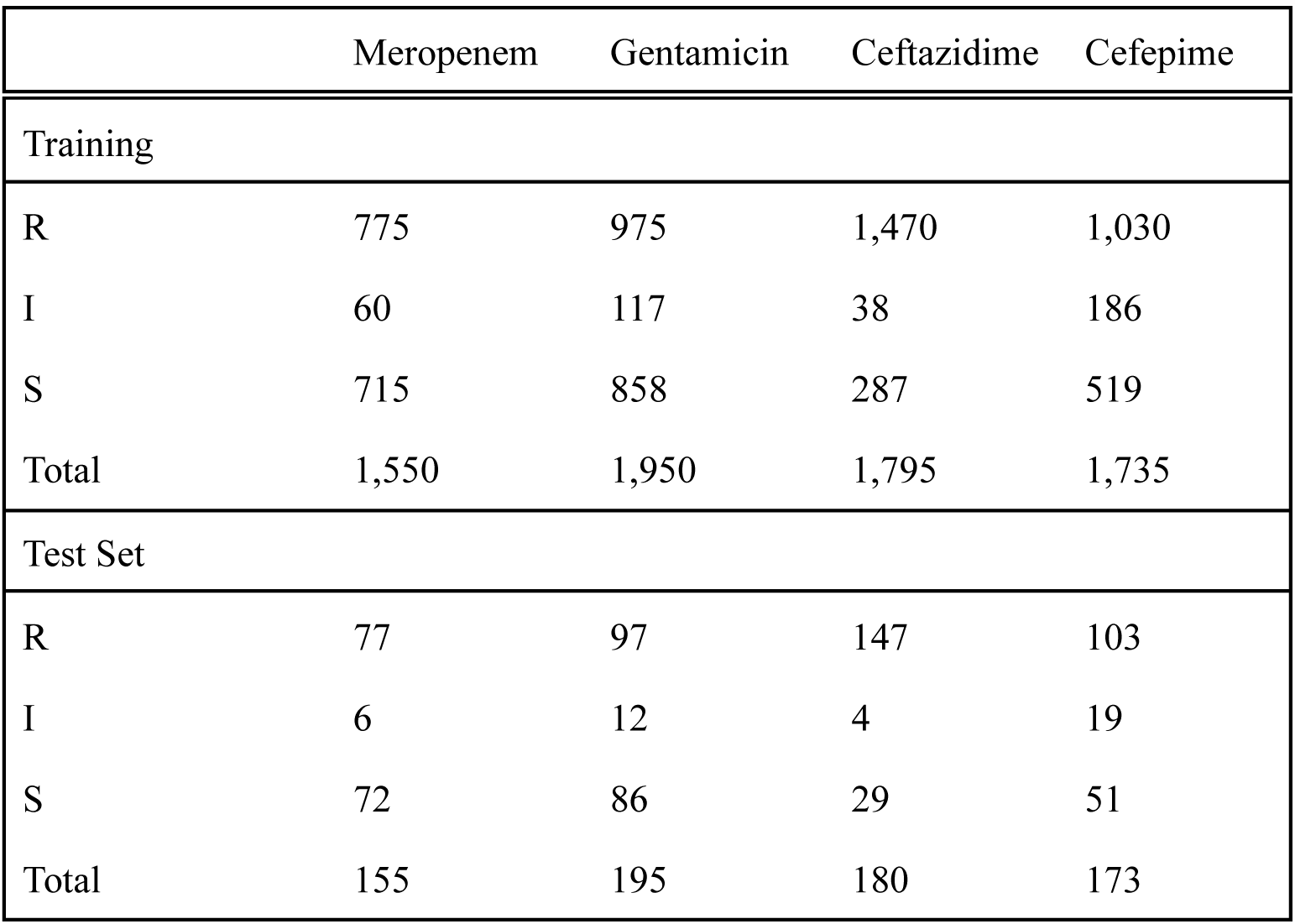
Datasets (training and test sets) for each antimicrobial agent. R: resistant sample; I: intermediate sample; S: susceptible sample.

### HYPERPARAMETER SPACE EXPLORATION

Hyperparameter optimization was conducted independently for each antimicrobial-specific model to account for differences in resistance prevalence, genomic signal complexity, and class imbalance. Although the underlying model architecture remained unchanged across all antimicrobials, the hyperparameter search was rerun to avoid implicit parameter transfer between antimicrobial contexts. The hyperparameters explored are in Table 2. Therefore, for each antibiotic, we trained 27 models with different hyperparameters, and we selected the best among the 27. The final hyperparameter configuration for each model was selected based on the best performance observed during the training/validation phase, ensuring an independent optimization process for FEP, CAZ, GEN, and MEM. For CAZ and FEP, due to class imbalance, the positive class weight (resistant) was reduced because the number of resistant samples exceeded the number of susceptible samples. The adjusted weight was calculated as the sum of susceptible (S) and intermediate (I) samples divided by the total number of resistant (R) samples. Table 3 reports the resulting weights for each antimicrobial agent.

**Table 2.**
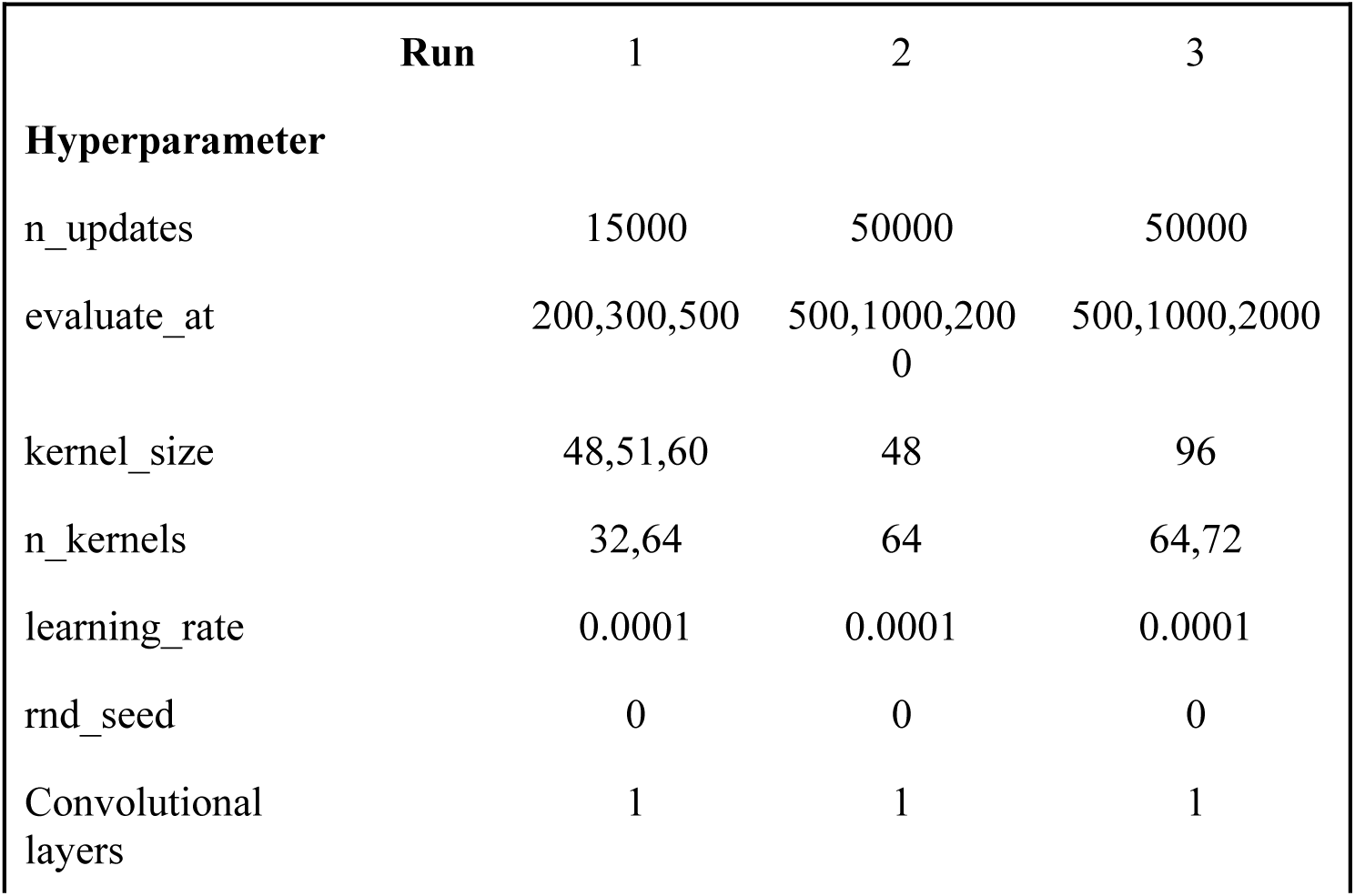
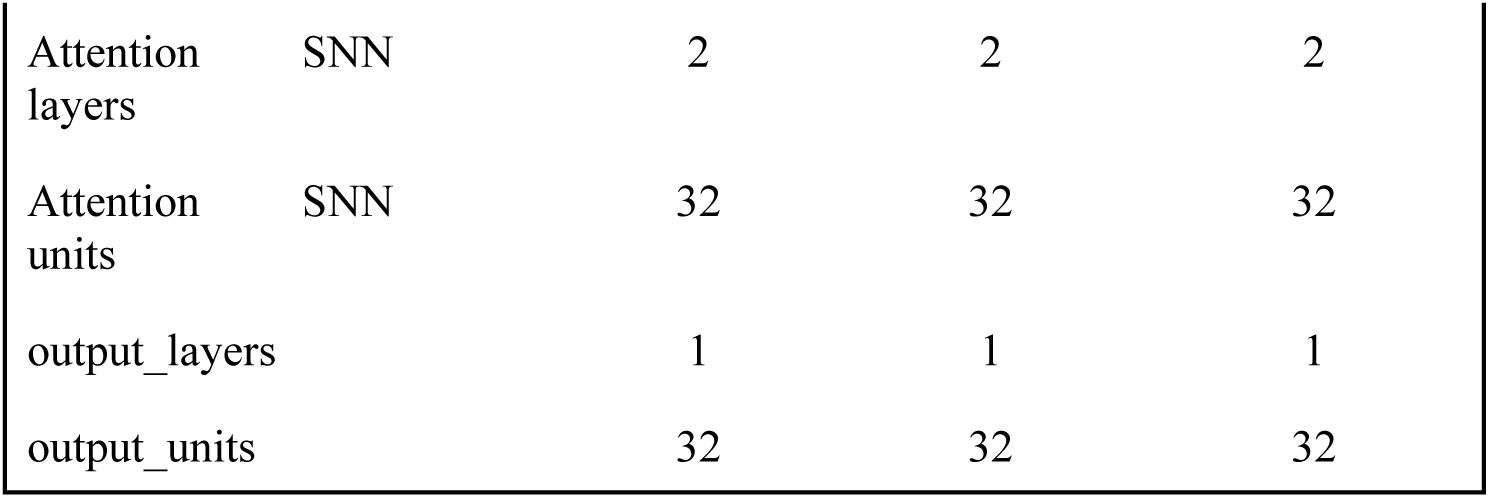
Hyperparameter space search used to generate the models. n_updates: corresponds to the total number of parameter updates during training; evaluate_at: indicates the evaluation frequency on the validation set; kernel_size: denotes the size of the convolutional filters applied to the sequences; n_kernels: represents the number of convolutional filters; learning_rate: is the learning rate used by the optimizer; rnd_seed: sets the random seed for reproducibility; Convolutional layers is the number of layers of the convolutional (embedding) network; attention SNN layers: defines the number of layers in the attention module; attention SNN units: corresponds to the number of internal units in the attention mechanism; output_layers: indicates the number of layers in the output network; output_units: represents the number of units in these layers; SNN stands for self-normalizing neural network (Klambauer et al., 2017), part of the Hopfield layer. Each run comprises multiple executions with each set of parameters listed.

| Run | 1 | 2 | 3 |
| --- | --- | --- | --- |
| **Hyperparameter** |  |  |  |
| `n_updates` | 15000 | 50000 | 50000 |
| `evaluate_at` | 200,300,500 | 500,1000,2000 | 500,1000,2000 |
| `kernel_size` | 48,51,60 | 48 | 96 |
| `n_kernels` | 32,64 | 64 | 64,72 |
| `learning_rate` | 0.0001 | 0.0001 | 0.0001 |
| `rnd_seed` | 0 | 0 | 0 |
| `Convolutional layers` | 1 | 1 | 1 |

|  |  |  |  |  |
| --- | --- | --- | --- | --- |
| Attention layers | SNN | 2 | 2 | 2 |
| Attention units | SNN | 32 | 32 | 32 |
| output_layers |  | 1 | 1 | 1 |
| output_units |  | 32 | 32 | 32 |

**Table 3.** Positive weights used on training for each antimicrobial.

|  | MEM | GEN | CAZ | FEP |
| --- | --- | --- | --- | --- |
| **Positive weight (pos_weight)** | 1 | 1 | 0.22 | 0.68 |

We used the Python Hydra library (https://hydra.cc/) to explore optimal parameter settings. After several training runs, we defined a standard set of hyperparameters for model generation (Table 4).

**Table 4.** Final hyperparameter configurations selected for each antimicrobial model.

| **Antimicrobial** | Meropenem | Gentamicin | Ceftazidime | Cefepime |
| --- | --- | --- | --- | --- |
| **Hyperparameter** |  |  |  |  |
| pos_weight | 1 | 1 | 0.22 | 0.68 |
| n_updates | 15000 | 50000 | 15000 | 15000 |
| evaluate_at | 500 | 2000 | 500 | 500 |
| kernel_size | 48 | 96 | 51 | 51 |
| n_kernels | 32 | 64 | 64 | 64 |
| learning_rate | 0.0001 | 0.0001 | 0.0001 | 0.0001 |
| sequence_embedding_layers | 1 | 1 | 1 | 1 |
| attention_layers | 2 | 2 | 2 | 2 |
| attention_units | 32 | 32 | 32 | 32 |
| output_layers | 1 | 1 | 1 | 1 |
| output_units | 32 | 32 | 32 | 32 |

### MODEL TRAINING

For deep learning model training, isolates categorized as intermediate (I) were grouped with susceptible (S) isolates (CLSI M100 2025; EUCAST 2025). The model architecture follows the DeepMDC framework and is inspired by DeepRC (Widrich et al., 2020), combining convolutional sequence encoders with a Modern Hopfield Network. The whole genome was annotated to identify open reading frames (ORFs), and only the resulting amino acid sequences were used and provided to the model as a bag of variable-length inputs. Isolates with classifier outputs greater than 0.5 were assigned to the resistant class (R), whereas outputs equal to or below this threshold were classified as susceptible (S). During training, we applied regularization via periodic validation at predefined evaluation steps, using a five-fold cross-validation procedure. The model state that yielded the lowest validation loss was saved and subsequently used to evaluate on the test set.

The classification network translates the representations into class probabilities (R, S). For training, we employed the Cross-Entropy Loss function and the Adam optimizer (Kingma & Ba, 2014) to update the model weights across the entire network. The selected hyperparameters for each antimicrobial model are summarized in Table 4.

For model training and inference, we used a server with the following configuration: 2x AMD EPYC 7543 2.8 GHz 32-core processors (64 physical cores total), 256 GB RAM, 32 TB HDD, 480GB SSD, and an NVIDIA A30 24GB GPU. We also used Amazon Web Services (AWS) *p5.4xlarge* instances with the following configuration: an AMD EPYC 7R13 processor with 8 physical cores, 16 vCPUs, 256 GB of RAM, and an NVIDIA H100 80GB GPU. EVALUATION METRICS: Evaluation metrics are important indicators of how well trained models perform on classification tasks. In this work, we have calculated four metrics: Receiver Operating Characteristic Area Under the Curve (ROC-AUC), balanced accuracy, F1 score, and Matthews Correlation Coefficient (MCC). We describe these metrics below:

**1- Balanced accuracy:** Balanced accuracy is a metric that corrects for the misleading results that standard accuracy can produce when evaluating machine learning models on imbalanced datasets. Instead of simply calculating the percentage of correct predictions, balanced accuracy ensures each class contributes equally to the final score. In binary classification, balanced accuracy is the average recall across classes. Recall for a class is also known as its true positive rate or sensitivity.

- *Sensitivity* (True Positive Rate): The percentage of positive cases correctly identified.

- *Specificity* (True Negative Rate): The percentage of negative cases correctly identified.

The formula is:

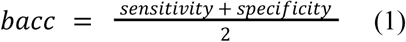

**2- Precision**: Precision is the proportion of correct positive predictions with respect to all (true or false) positive predictions returned by the model:

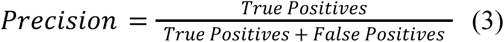

**3- Recall**: Recall (or True Positive rate) is the proportion of actual positive measures correctly predicted by the model:

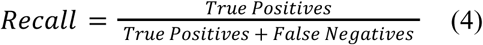

**4- F1 score:** F1 score is the harmonic mean of Precision and Recall, and is a single parameter that balances these other two metrics:

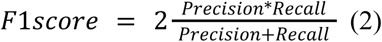

The highest possible value of the F1 score is 1.0, indicating perfect precision and recall, and the lowest possible value is 0.

**5- Receiver Operating Characteristic Area Under the Curve (ROC-AUC):** Receiver Operating Characteristic (ROC) Area Under the Curve (AUC) is a performance metric that summarizes a binary classifier’s ability to distinguish between positive and negative classes. It is a single scalar value ranging from 0.5 to 1.0, where 0.5 indicates no discriminative ability (equivalent to random guessing), and 1.0 signifies a perfect classifier. The AUC is interpreted as the probability that a randomly selected positive sample is ranked higher than a randomly selected negative sample. It is calculated by integrating the ROC curve. The ROC curve is created by plotting the True Positive Rate against the False Positive Rate at various threshold settings.

**6- Matthews Correlation Coefficient (MCC):** MCC is a metric that evaluates the quality of binary classification models, providing a balanced measure even for imbalanced datasets. Ranging from -1 to +1, the MCC considers all four outcomes from a confusion matrix (True Positives, True Negatives, False Positives, False Negatives). A score of +1 indicates a perfect prediction, 0 suggests random guessing, and -1 signifies a complete disagreement between prediction and actual values. The formula for the Matthews Correlation Coefficient (MCC) for binary classification derives from the four values in a confusion matrix: true positives (TP), true

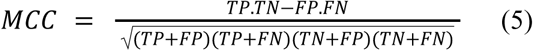

### ATTENTION COEFFICIENTS RETRIEVAL

For attention-weight retrieval after an inference, we used the scripts *extract_attention_weights.py* and *prediction_and_att_weights.py*, available in the companion repository (https://github.com/CABGenOrg/deepmdc). Those scripts extract the attention coefficients for each ORF in each sample. The *extract_attention_weights.py* script extracts all coefficients, resulting in extensive tables, while the *prediction_and_att_weights.py* script collects only the top 5 attention coefficients for each sample. We noted that the attention coefficients of a sample tend to zero very quickly after the fifth-largest attention coefficient. The *prediction_and_att_weights.py* script also displays the model’s prediction for the corresponding sample and whether the prediction is correct. The execution of either script follows the same parameters as the test script.

### DATA ANALYSIS

We used Python version 3.11, along with its modules matplotlib (version 3.10.6), NumPy (version 2.3.2), pandas (version 2.3.2), and seaborn (version 0.13.2) to analyze the data. The analysis script, *data_analysis.py*, is available in the companion repository (https://github.com/CABGenOrg/deepmdc). We divided the analysis into two parts: attention coefficient frequency analysis and performance metrics. Initially, we merged the datasets from the five folds into a single dataset, and an additional column, “model,” was added to specify the originating model for each record. We identified each amino acid sequence as a unique product and divided the samples into two groups based on their combination of predicted and labeled classes. Our first group includes samples predicted and labeled as sensitive (true negatives), while our second group includes those with a resistance match (true positives). The frequency analysis aimed to compute how often each product appeared in the best model for each antibiotic. We analyze these frequencies within each group separately, computing the top 10 for each group and antibiotic for better readability. Finally, we generated performance metric graphics using information from the validation and test stages, presenting the metrics described earlier.

## RESULTS

### DEEPMDC - INTERPRETABLE DEEP LEARNING ARCHITECTURE FOR BACTERIAL GENOME PROFILING

We adapted the DeepRC architecture (Widrich et al., 2020) to process bacterial genomes. In the **first stage**, we divide the entire genome into ORFs, which we then translate into amino acid sequences. In this architecture, all ORF instances compose the DeepMDC input, including small ORFs (sORF). In the **second stage**, a predefined encoder processes the bag of ORFs and produces a latent-space representation (Figure 1). The encoder can be a 1D convolutional network (CNN), an LSTM, or a Genomic Language Model (Zhou et al., 2023; Dalla-Torre et al., 2025). The output of the encoder is fixed-sized representations for every ORF. The 1D CNN output vector size equals the predefined number of convolutional kernels, which may vary by antibiotic (see Table 4).

**Figure 1.**
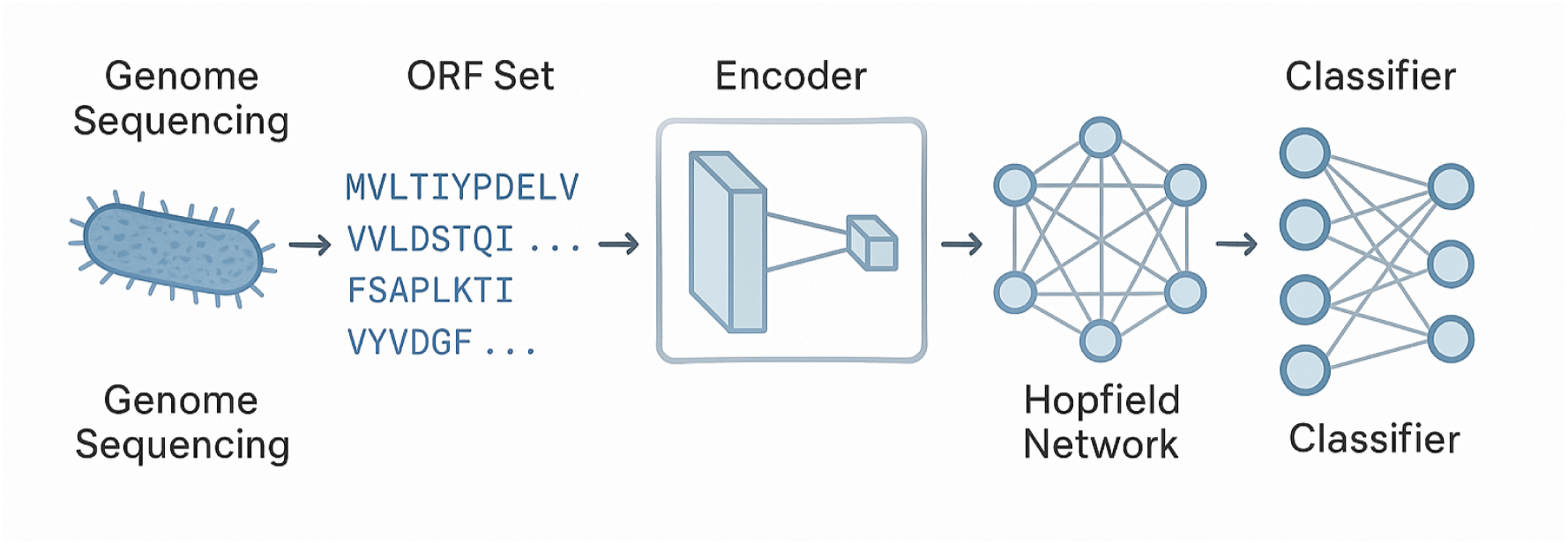
DeepMDC architecture for processing bacterial whole-genome sequences. We evaluated this architecture to classify bacterial strains into antimicrobial-susceptible/resistant phenotypes. We treated intermediate strains as susceptible, yielding a binary classifier. In this work, the encoder was a 1D convolutional neural network (CNN). A 1D CNN is more computationally efficient than transformer-based or LSTM encoders and requires less VRAM on a GPU for training and inference. However, it yields good results, as shown below.

In the 1D CNN, each filter (or kernel) is responsible for detecting subsequences of length k, the kernel size. This operation evolves each kernel to detect biological motifs that help distinguish between antimicrobial resistance classes (in this work, either resistant or sensitive). 1D CNN processing converts each bacterial ORF into a fixed-length feature vector whose length corresponds to the number of kernels. For each position of this vector, the convolutional layer performs global max-pooling over all ORF positions relative to the associated kernel. In Table 4, we can observe several differences among the antibiotics in the number of kernels and kernel size of the 1D convolutional networks. These differences reflect different action mechanisms. For instance, both ceftazidime and cefepime are cephalosporins and have the same number of kernels and kernel size. Gentamicin is a powerful aminoglycoside, and meropenem is a broad-spectrum carbapenem antibiotic; each has its own values for the number of kernels and kernel sizes, suggesting that the 1D convolutional network should be tailored to antibiotic classes.

In the **third stage**, a dense associative memory layer (Modern Hopfield Network) processes the fixed-size latent representations via a self-normalizing neural network (Klambauer et al., 2017) and a softmax function to assign an attention coefficient to each ORF representation, since the update rule of the Modern Hopfield Network is equivalent to the Transformer’s self-attention mechanism (Ramsauer et al., 2020). The Modern Hopfield Network then generates a fixed-length vector (its size equals the number of convolutional kernels) via attention pooling, which serves as input to an MLP classifier. This operation is described by equation 6.

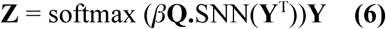

Where **Z** is the output of the Modern Hopfield Network, **Q** is a learned query vector and represents the summarized resistance profile of a species to an antibiotic, and **Y** are the 1D CNN encoded representations of the ORFs. The keys (i.e., the stored patterns) of the self-attention mechanism are generated by a self-normalizing neural network (SNN) that has the ORF representations as input. SNNs are deep feed-forward networks that automatically keep their layer activations at zero mean and unit variance, eliminating the need for explicit normalization techniques, and have been successfully applied to process genomic data (Li et al., 2020). In DeepMDC, the SNN processes the encoding matrix **Y**, thereby outputting the stored patterns in the Hopfield Network, which serve as the keys of the self-attention mechanism. The resulting vector **Z** is then processed by a fully connected network for final classification.

In this work, we initially implemented a single attention head and utilized attention coefficients assigned to the ORFs during inference to interpret the model’s output. Therefore, the higher the attention weight attributed to an ORF, the higher the importance of that ORF in the inference decision process. It is worth noting that the sum of attention weights over all ORFs of a bacterial genome (> 5000 ORFs) is equal to one due to the softmax function. In practice, we observe that attention weights tend to zero rapidly after the five highest weights for a genome, due to the SNN used to compute attention coefficients. For this reason, we focus on the five highest-attention-weighted ORFs for each genome below.

In summary, the CNN learns to recognize patterns in ORFs; the modern Hopfield network identifies and prioritizes the most relevant ORFs; the classification network assesses whether the prediction approximates the actual label. We fine-tuned all networks jointly to improve performance.

### DEEPMDC ACHIEVES SOLID PERFORMANCE METRICS FOR SEVERAL ANTIBIOTICS

#### Meropenem

The validation results for each of the five folds are in Figure 2a. One can observe that folds 1 and 2 achieved area under the curve, balanced accuracy, and F1 scores of 0.9 or higher. MCC in these two models is 0.8 or better.

**Figure 2.**
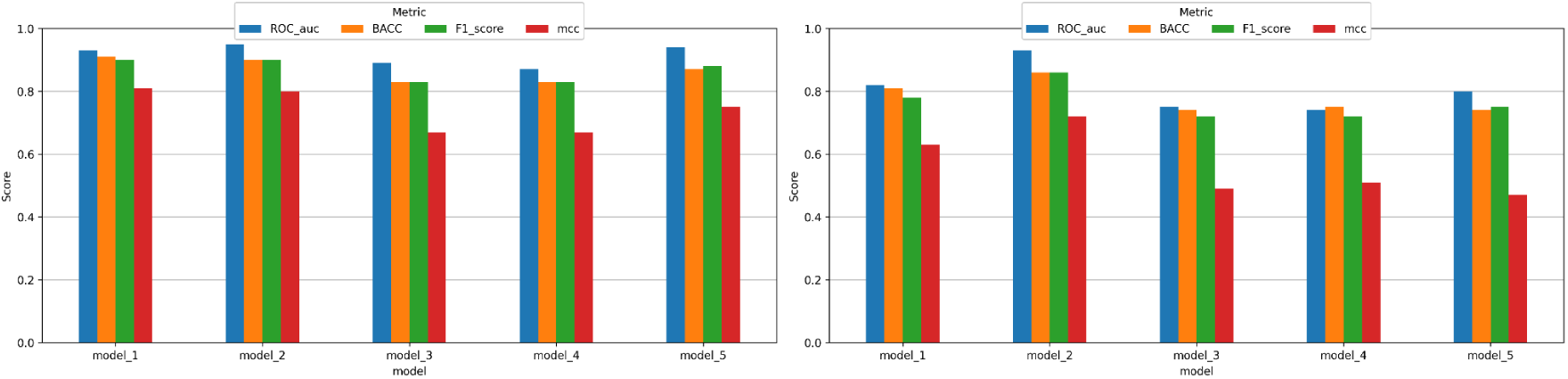
(a) Performance metrics in validation stage for the meropenem model; (b) Performance metrics from test dataset for the meropenem model.

For testing DeepMDC, we treated each fold generated during training/validation as a separate model. We observed that fold two (also referred to in this work as model 2) stands out with a Receiver Operating Characteristic AUC of 0.93, a balanced accuracy of 0.86, a F1 score of 0.86, and an MCC of 0.72 (Figure 2b). In this case, the best model assigned for meropenem was model (fold) 2. Complete results and statistics for meropenem are in tables S1 and S2.

We separated the test set samples from the best model into two subgroups: S-S (evaluated as susceptible by both DeepMDC and AST) and R-R (evaluated as resistant by both DeepMDC and AST). Figure 3 shows the proteins that appear most frequently in the five highest attention weights set for correctly classified samples in both the S-S (Figure 3a) and R-R (Figure 3b) groups in the best performance model (model 2). Several ORFs relevant to *K. pneumoniae* resistance stand out in the R-R group, including the Tn3-like element Tn*4401* family transposase. For the S-S group, the set of ORFs with the highest attention weights differs entirely. Some proteins associated with susceptibility include *OmpK36* and the associated outer membrane porin (Figure 3).

**Figure 3.**
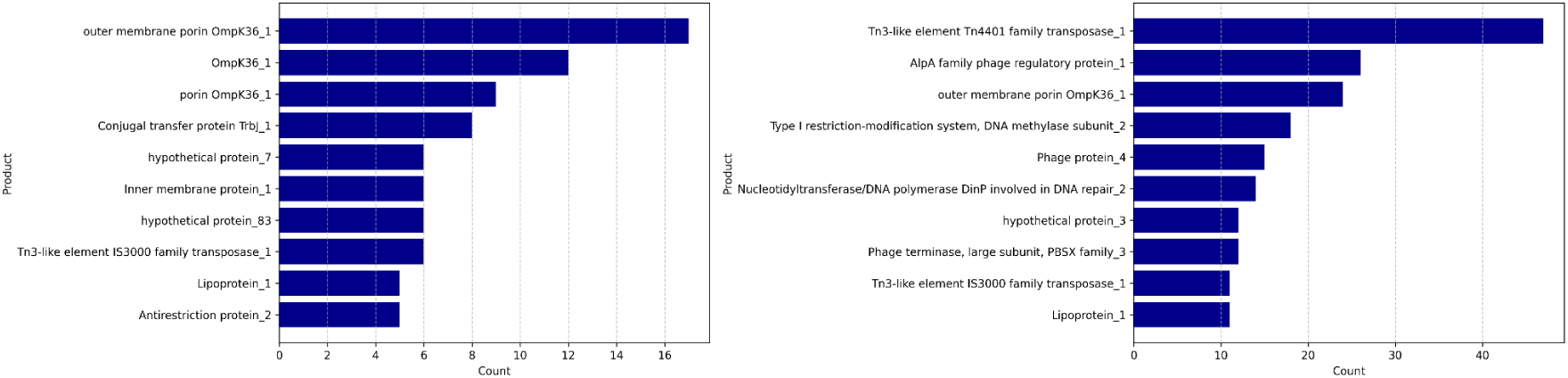
(a) Highest attention coefficients for the meropenem antibiotic in model 2 within group S-S; (b) Highest attention coefficients for the meropenem antibiotic in model 2 within group R-R.

The identification of these elements suggests that DeepMDC captures both canonical resistance determinants and broader genomic contexts associated with antimicrobial susceptibility. Transposable elements, such as those from the Tn3 family, are known to mediate the dissemination of resistance genes across strains and species (Cuzon et al., 2011), while porins such as OmpK36 are directly involved in antibiotic influx and have been repeatedly associated with altered susceptibility profiles (Sugawara et al., 2016). This indicates that the model does not rely on isolated genetic markers but instead integrates signals from multiple genomic features that collectively contribute to the resistance phenotype.

#### Cefepime

Model 1 demonstrated the best performance across the five evaluated folds and was therefore selected for subsequent analyses (Figure 4). For this model, ROC-AUC was 0.79, balanced accuracy was 0.71, F1 score was 0.74, and MCC was 0.42 in the validation process, with ROC-AUC of 0.83, balanced accuracy of 0.76, F1 score of 0.79, and MCC of 0.51 on the test set. Complete results and statistics for cefepime are in tables S3 and S4.

**Figure 4.**
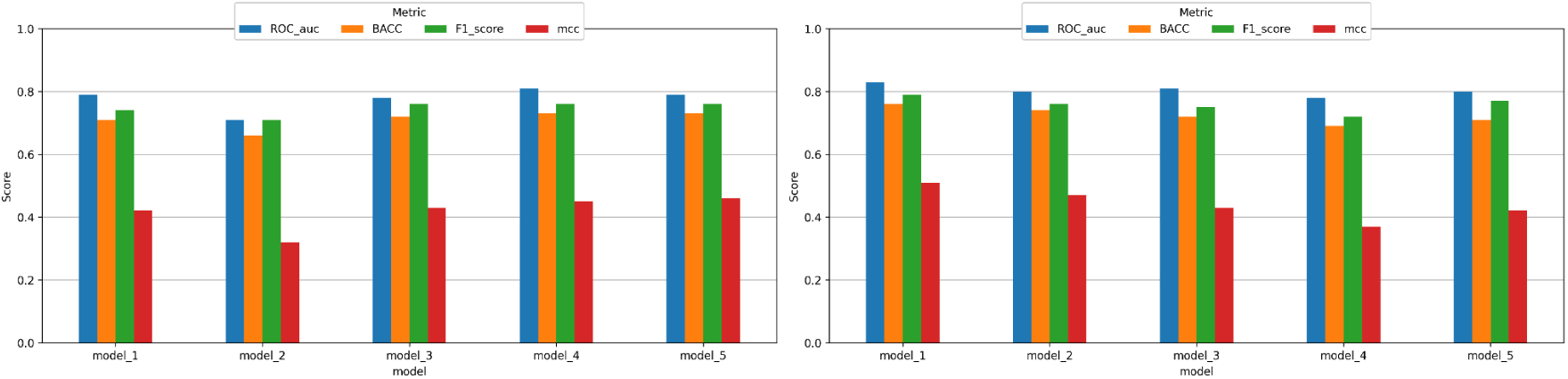
(a) Performance metrics in validation stage for the cefepime model; (b) Performance metrics from test dataset for the cefepime model.

Among the 5 most frequent ORFs with high attention observed in the R-R group are the streptomycin 3’’-adenylyltransferase, the *YicG* protein, and *ninF* (lambda phage). For the S-S group, the NADH dehydrogenase and the *YpdK* membrane protein stand out. (Figure 5).

**Figure 5.**
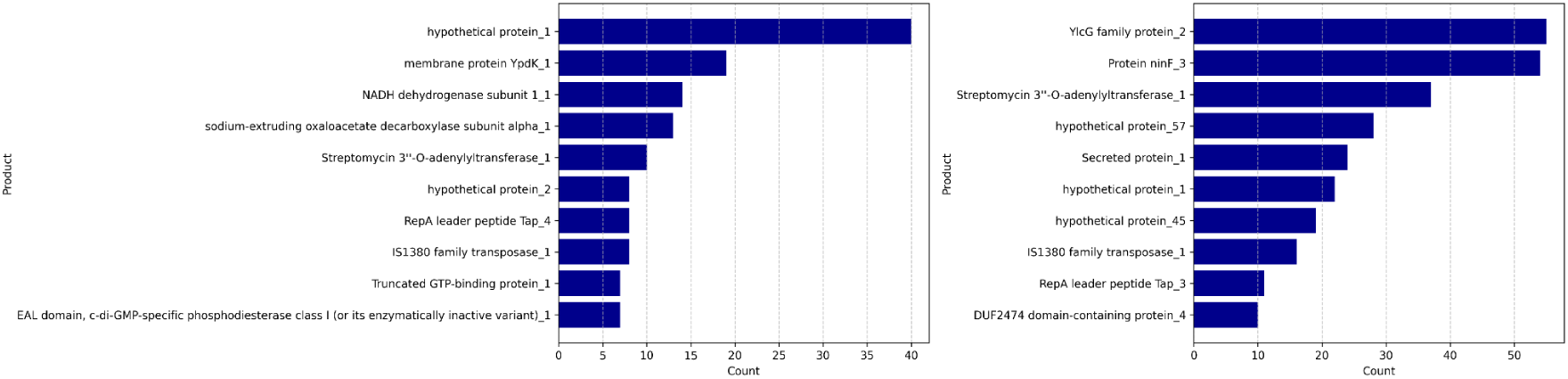
(a) Highest attention coefficients for the cefepime antibiotic in model 1 within group S-S; (b) Highest attention coefficients for the cefepime antibiotic in model 1 within group R-R.

#### Ceftazidime

For ceftazidime, Model 1 exhibited the best performance (Figure 6). For model 1, during validation, ROC-AUC was 0.94, F1 score was 0.92, MCC was 0.66, and balanced accuracy was 0.87. For the test set, the model has an ROC-AUC of 0.88, an F1 score of 0.94, an MCC of 0.72, and a balanced accuracy of 0.88. Beyond these strong performance results, it is worth noting that ceftazidime exhibits the greatest training-data imbalance among the antibiotics considered. Complete results and statistics for ceftazidime are in tables S5 and S6.

**Figure 6.**
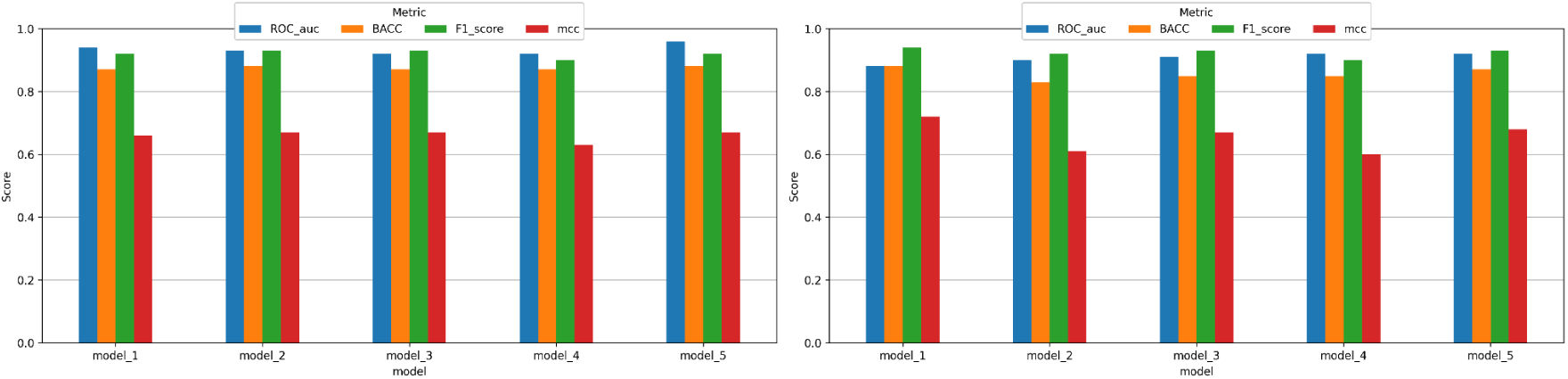
(a) Performance metrics in validation stage for the ceftazidime model; (b) Performance metrics from test dataset for the ceftazidime model.

Among the most frequently identified high-attention ORFs in the R-R group is a mutated variant of the Colicin V protein (Figure 7b). The Colicin V protein is also present in the S-S group, but with a different mutation than that observed in the R-R group (Figure 7a).

**Figure 7.**
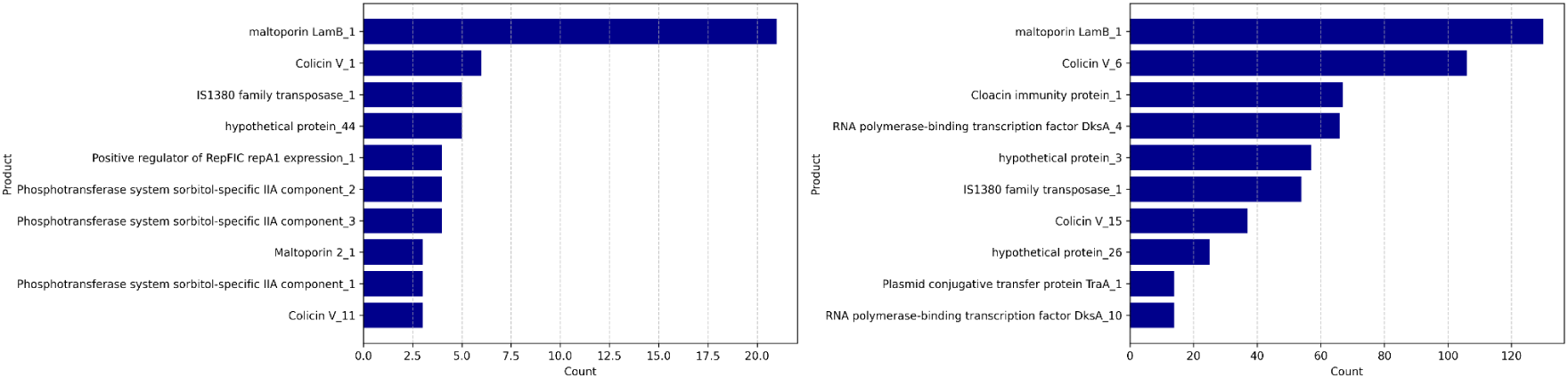
(a) Highest attention coefficients for the ceftazidime antibiotic in model 1 within group S-S; (b) Highest attention coefficients for the ceftazidime antibiotic in model 1 within group R-R.

#### Gentamicin

For gentamicin, Model 1 exhibited the best performance and was selected for downstream analyses (Figure 8). The balanced accuracy for model 1 during validation was 0.89, with an ROC-AUC of 0.94, an F1 score of 0.89, and an MCC of 0.78. On the test set, model 1 achieved an ROC-AUC of 0.89, a balanced accuracy of 0.81, an F1 score of 0.81, and an MCC of 0.62. Complete results and statistics for gentamicin are in tables S7 and S8.

**Figure 8.**
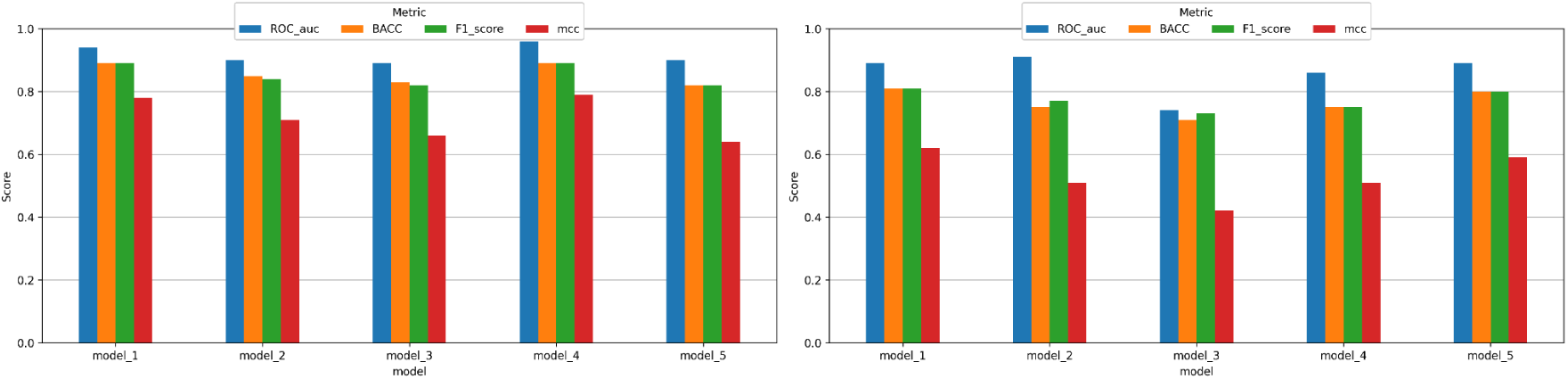
(a) Performance metrics in validation stage for the gentamicin model; (b) Performance metrics from test dataset for the gentamicin model.

For the gentamicin R-R group, we highlight the IS3 family ISKpn11 transposase (Figure 9b). For the S-S group, frequent high-attention proteins are holin and a Na+ symporter (Figure 9a).

**Figure 9.**
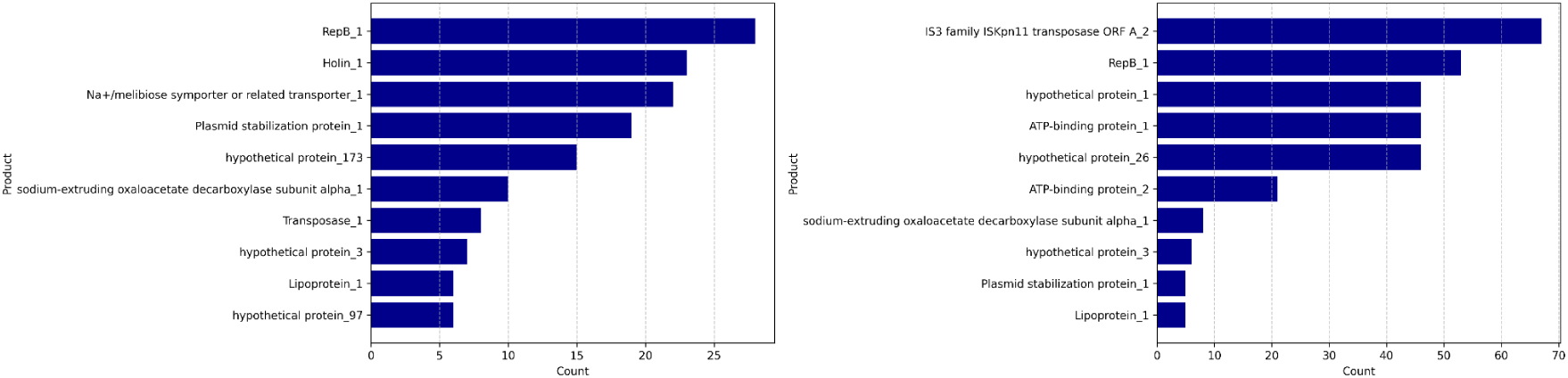
(a) Highest attention coefficients for the gentamicin antibiotic in model 1 within group S-S; (b) Highest attention coefficients for the gentamicin antibiotic in model 1 within group R-R.

## DISCUSSION

In this study, we applied deep learning models to *K. pneumoniae* genomic and proteomic data to predict resistance phenotypes to cefepime, ceftazidime, gentamicin and meropenem to identify protein features associated with these predictions. The results obtained with DeepMDC demonstrate that the model not only achieves high predictive performance but also captures biologically meaningful genomic patterns underlying AMR in *K. pneumoniae*. In contrast to rule-based approaches, such as ResFinder and PointFinder, which rely on curated databases linking specific genes and mutations to resistance phenotypes, DeepMDC adopts a fundamentally different strategy. It learns discriminative representations directly from raw sequence data, without requiring prior identification of elements associated with resistance. This design allows DeepMDC to capture signals not only from well-characterized resistance genes but also from previously unannotated or poorly understood genomic regions. The classical machine learning approaches, while often achieving competitive predictive performance, typically rely on predefined feature sets, such as gene presence/absence matrices derived from curated databases. As a result, their predictive capacity is inherently constrained by their knowledge of resistance mechanisms, limiting their ability to uncover novel genomic determinants (Aytan-Aktug et al., 2020; Peng et al., 2022).

The attention-based interpretability analysis of our deep learning model provided critical insights into the genomic signals that influence its predictions, highlighting both the biological coherence and the interpretive limitations of antimicrobial resistance (AMR) prediction. By examining the distribution of attention coefficients across gene products, we identified how the model differentially prioritized classical resistance determinants versus regulatory or structural features, particularly in misclassified isolates. Overall, the model prioritized biologically meaningful genomic features commonly associated with antimicrobial resistance. Interestingly, several hypothetical proteins also exhibited high attention weights, suggesting potential roles of uncharacterized loci co-localized with resistance determinants. Identifying the functions of the hypothetical proteins will contribute to a better understanding of the bacterial proteome and its role in virulence (Dey et al., 2022). In this regard, DeepMDC can function as a prioritization tool, highlighting hypothetical proteins that consistently receive high attention weights across independent training folds as candidates for functional characterization. The biological relevance of these candidates may be further supported by their genomic context, particularly when they occur with known resistance determinants. Rather than indicating direct causal involvement, these attention signals are more likely capturing broader genomic contexts statistically associated with resistance, such as conserved plasmid architectures or insertion sequence neighborhoods (Partridge et al., 2020; Rodríguez-Beltrán et al., 2021). Nevertheless, the recurrent identification of hypothetical proteins across independent models provides a rational and data-driven starting point for experimental validation, potentially contributing to the discovery of novel loci associated with resistance in *K.pneumoniae*.

We displayed the 10 most frequent proteins with high attention coefficients across the 5-fold cross-validation models, highlighting elements strongly associated with antimicrobial resistance and genomic plasticity in *K. pneumoniae*. To explore the biological relevance of the model’s predictions, we analyzed the distributions of attention coefficients for the most frequent predicted products across two groups (S-S and R-R) and different antimicrobials (cefepime, ceftazidime, gentamicin, and meropenem). The use of attention mechanisms as an interpretability layer in genomic deep learning models enables the identification of biologically relevant features associated with antimicrobial resistance (Hu et al., 2024; Su et al., 2019).

An important observation is the variation in model performance both across antimicrobials and between validation and tests stages. Among the four models, meropenem achieved the highest discriminative performance (ROC-AUC 0.93), followed by gentamicin and ceftazidime (ROC-AUC 0.89 and 0.88, respectively), while cefepime presented the lowest performance (ROC-AUC 0.83). The comparatively lower performance of the cefepime model may reflect the greater mechanistic heterogeneity associated with resistance to fourth-generation cephalosporins, which can involve multiple partially overlapping mechanisms including ESBL production, AmpC overexpression, and porin loss (Wyres *et al*., 2020; Hu *et al*., 2024).

Notably, the cefepime model showed an improvement from validation to test, which, although uncommon, may suggest that the validation splits contained a more challenging distribution of samples, such as borderline or noisy phenotypes. In contrast, ceftazidime showed a modest performance decrease from validation to test, consistent with expected generalization behavior. Together these results highlight that the model performance in genomic AMR prediction is strongly dependent on the underlying biological complexity of each drug class and emphasize the importance of reporting both validation and test metrics separately (Hu *et al*., 2024; Aytan-Aktug *et al*., 2020).

For instance, for meropenem, our findings reveal that the models captured a multi-layered genomic signature, integrating mobility, envelope remodeling, and regulatory systems consistent with the evolutionary strategies underlying the dissemination of carbapenem resistance in Enterobacterales such as *K. pneumoniae* (Partridge et al., 2018; Wyres et al., 2018; Rodríguez-Beltrán et al., 2021; Lam et al., 2021).

In the meropenem R-R group (predicted resistant, actually resistant), DeepMDC assigned high-attention weights to Tn*4401*-like transposases (also relevant in the gentamicin model), *OmpK36* porins, and phage regulatory proteins (also present in the cefepime model). These elements are known to be involved in the mobilization of the carbapenemase gene (*bla*_KPC_) and in porin modification, both of which are critical for carbapenem resistance (Doménech-Sánchez et al., 1999; Hernández-Allés et al., 1999; Sugawara et al., 2016).

Based on these findings, the transposon Tn*4401* emerges as a central player in the dissemination of the *bla*_KPC_ gene. This Tn*3*-family transposon plays a pivotal role in disseminating the *bla*_KPC_ gene, which encodes a class A carbapenemase that hydrolyzes penicillins, cephalosporins, and carbapenems (Naas et al., 2008; Cuzon et al., 2011). Several isoforms (e.g., Tn*4401a*, Tn*4401b*, Tn*4401c*) have been described, differing by small deletions or rearrangements in the promoter region upstream of *bla_KPC_*, which can affect expression levels and resistance phenotypes (Naas et al., 2008; Cuzon et al., 2011). Frequently carried on conjugative plasmids of different incompatibility groups (IncFII, IncN, IncL/M), Tn*4401* facilitates horizontal transfer among Enterobacterales, driving the global spread of carbapenem resistance (Mathers et al., 2015; Chen et al., 2014). Consequently, Tn*4401* is a key mobile genetic element that underpins the epidemic dissemination of KPC-producing *K. pneumoniae* and other multidrug-resistant *Enterobacterales*.

From a microbiological perspective, these findings reinforce that carbapenem resistance is primarily driven by horizontal gene transfer and plasmid epidemiology, rather than de novo mutation alone. Population genomic studies have shown that epidemic clones of *K. pneumoniae* act as hubs for resistance gene dissemination, frequently carrying Tn4401-associated plasmids (Wyres et al., 2020; Partridge et al., 2018). The identification of transposases in multiple antibiotic models (including gentamicin) is consistent with the fact that mobile genetic elements often carry clusters of resistance genes, leading to co-selection under antibiotic pressure (Davies & Davies, 2010; Botelho & Schulenburg, 2021). Thus, the model appears to capture genomic signatures of resistance at the level of mobile resistance networks, rather than isolated genes.

Beyond gene mobilization, outer membrane permeability barriers also contribute substantially to the carbapenem resistance phenotype. The strong attention assigned to OmpK36 highlights the importance of membrane permeability in β-lactam resistance. OmpK36 is one of the main porins responsible for antibiotic influx in *K. pneumoniae*, particularly for carbapenems (Sugawara et al., 2016; Tsai et al., 2011). Alterations in OmpK36 (including reduced expression, truncation, or structural mutations) significantly reduce antibiotic penetration and are strongly associated with carbapenem resistance (David et al., 2022; Wong et al., 2019). Importantly, porin loss alone is often insufficient to confer high-level resistance but acts synergistically with β-lactamases, particularly KPC and OXA-type enzymes (Meekes et al., 2025; Wong et al., 2022). Conversely, the presence of intact porins in susceptible isolates reflects preserved permeability, which is a well-established determinant of β-lactam susceptibility (Hernández-Allés et al., 1999; Doménech-Sánchez et al., 1999). These findings suggest that the model distinguishes functional states of membrane transport systems, rather than simply detecting gene presence.

The focus on DNA methylases also suggests that the model also captured epigenetic or regulatory elements relevant to horizontal gene transfer. DNA methylation has been shown to modulate gene expression, replication, and conjugative transfer of mobile genetic elements, thereby contributing to the dissemination of antimicrobial resistance determinants (Roberts et al., 2015; Oliveira et al., 2020). Emerging evidence indicates that epigenetic mechanisms can modulate plasmid stability and expression of resistance genes, contributing to adaptive responses under antibiotic pressure (Casadesús & Low, 2006). Therefore, the model’s sensitivity to these features suggests that it captures not only structural resistance determinants but also regulatory contexts that modulate resistance expression.

In the meropenem S–S Group (true sensitive isolates), attention was focused on porins (*OmpK36*), phage structural proteins, and transfer proteins (TrbJ), reflecting genomic stability and preserved permeability, typical of susceptible isolates lacking mobile resistance determinants (Davies & Davies, 2010; Shintani et al., 2015). The model’s close attention to porin (and other transport-related proteins in the cefepime, ceftazidime, and gentamicin models), likely intact in sensitive isolates, indicates that it correctly recognized the preservation of membrane permeability as a hallmark of susceptibility. In contrast, resistant isolates typically exhibit disruptions, frameshift mutations, or downregulation of *OmpK36*, resulting in decreased carbapenem influx (Strahilevitz et al., 2024; Palomba et al., 2025).

The model also emphasized phage proteins and AlpA-like regulators, which reflect the presence of stable prophage remnants or non-lytic phage elements integrated into the bacterial chromosome. In susceptible isolates, these phage-associated genes often play neutral or regulatory roles rather than contributing directly to resistance (Touchon et al., 2017; Colavecchio et al., 2017). In contrast, in resistant isolates, phage-related elements are frequently found adjacent to or within mobile genetic islands that carry antibiotic resistance genes (ARGs) (Partridge et al., 2018; Botelho & Schulenburg, 2021). In addition to regulatory mechanisms, the presence of phage-associated proteins among high-attention features reflects the role of bacteriophages in genomic plasticity. Temperate phages contribute to horizontal gene transfer and can facilitate the dissemination of resistance genes across bacterial populations (Touchon et al., 2017; Colavecchio et al., 2017). Phage elements are frequently integrated within genomic islands that harbor antimicrobial resistance genes, making them reliable indicators of genomic regions associated with resistance (Penadés et al., 2015). This suggests that the model detects broader genomic architecture linked to resistance, rather than individual loci alone.

Furthermore, the attention weight assigned to TrbJ, a component of the type IV secretion system, underscores its role in conjugation-dependent horizontal gene transfer. This level of attention weight may be unexpected in sensitive isolates. However, experimental studies show that Trb-like transfer systems are encoded by a wide variety of cryptic or non-conjugative plasmids, which may persist without actively mediating the dissemination of resistance (Shintani et al., 2015; Smillie et al., 2010; Gordils-Valentin et al., 2024).

In the R-R group, DeepMDC assigned high attention to streptomycin 3”-adenylyltransferase, an enzyme associated with aminoglycoside resistance across bacterial species. Although not directly related to β-lactam resistance, this observation may reflect co-selection processes, where resistance genes from different antibiotic classes are co-located on the same mobile genetic elements (Partridge *et al.,* 2018; Rodríguez-Beltrán *et al.,* 2021). Therefore, the model may be capturing genomic associations rather than direct mechanistic drivers of cefepime resistance. Additionally, the presence of LysR-type regulators such as *YicG* suggests that regulatory networks may contribute to adaptive responses in resistant strains, potentially through transcriptional reprogramming of stress response or efflux systems (Davies & Davies, 2010). The presence of phage-related proteins such as ninF further supports the role of mobile genetic elements in shaping these resistance-associated genomic contexts (Kumavath et al., 2025), consistent with findings from the meropenem model.

In the cefepime S-S group, the most attended features included metabolic and membrane-associated proteins, such as NADH dehydrogenase subunit 1 and *YpdK* (Figure 5a). This interpretation supports the idea that microbial susceptibility is not only the absence of resistance genes but also the preservation of metabolic balance (Davies & Davies, 2010).

For ceftazidime, DeepMDC achieved strong performance, despite a marked class imbalance in the dataset (Table 1). This result suggests that the model effectively captured consistent genomic signatures associated with resistance, likely reflecting the high prevalence of ESBL-producing strains among high-risk *K.pneumoniae* lineages (Wyres at al., 2020; Lam et al., 2021). Resistance to cefepime and ceftazidime is primarily mediated by ESBLs and AmpC β-lactamases (Paterson & Bonomo, 2005; Bush & Jacoby, 2010). The association with metabolic enzymes suggests that bacterial physiology influences antibiotic susceptibility. Metabolic activity has been shown to modulate antibiotic lethality through effects on reactive oxygen species and energy metabolism (Peng et al., 2015; Lobritz et al., 2015).

One notable finding was the high attention assigned to Colicin V in both resistant and susceptible groups, but with different mutational variants as shown in Figure 7. This distinction suggests that DeepMDC may be sensitive to fine-grained sequence variation in accessory genes, potentially reflecting differences in the genomic context of resistant versus susceptible isolates. Further work is needed to determine whether these Colicin V variants are linked to specific plasmid backbones or resistance-associated genetic modules.

For gentamicin, in the R-R group, the most prominent feature was the IS3 family transposon ISKpn11. This observation suggests that the model is capturing the mobilization context of resistance genes, rather than targeting specific enzymes that modify aminoglycoside directly. Insertion sequences such as ISKpn11 are known to facilitate gene mobilization and expression, and their presence may indicate genomic regions enriched in resistance determinants (Botelho & Schulenburg, 2021; Patridge et al, 2018). For gentamicin resistance, proteins such as ISKpn11 transposase were consistently prioritized. The identification of mobile genetic elements aligns with the known role of aminoglycoside-modifying enzymes, which are typically encoded on plasmids and transposons. Aminoglycoside resistance in *K. pneumoniae* is frequently mediated by aminoglycoside-modifying enzymes, including acetyltransferases, phosphotransferases, and adenylyltransferases (Ramirez, and Tolmasky, 2010; Garneau-Tsodikova and Labby, 2016). While not all highlighted proteins correspond directly to known modifying enzymes, their association with resistance phenotypes may reflect genomic contexts enriched for aminoglycoside resistance determinants or stress-response pathways activated under antibiotic pressure.

Additionally, the recurrent identification of hypothetical proteins suggests that the model captures previously uncharacterized genomic regions associated with resistance. Functional annotation studies have demonstrated that many hypothetical proteins are involved in virulence, stress response, and antimicrobial resistance (Dey et al., 2022; Galperin & Koonin, 2004). This highlights the potential of DeepMDC as a tool for discovering novel resistance-associated loci.

Across all four antimicrobials, a consistent pattern emerges in which DeepMDC prioritizes features associated with three broad biological dimensions: (i) mobile genetic elements, including transposases and phage-related proteins; (ii) membrane and transport systems, such as porins and transporters; and (iii) regulatory components, including transcription factors. This triad reflects the multi-layered nature of antimicrobial resistance, which arises from interplay between gene acquisition, cellular permeability, and regulatory adaptation (Ferrand et al., 2020). In this context, DeepMDC does not appear to rely on single markers, but rather captures distributed genomic signals that collectively define resistance phenotypes (Widrich et al., 2020; Ramsauer et al., 2020). Beyond the analysis of meropenem, the cross-antibiotic results suggest that DeepMDC can capture both shared and drug-specific genomic signals associated with antimicrobial resistance. We can improve the DeepMDC architecture in several ways. Replacing 1D convolutional networks in the encoder phase with LSTMs or transformer-based architectures warrants further investigation. For instance, we can replace the 1D CNN layer with a BERT-like encoder such as ESM-2 (Lin et al., 2023). However, we can expect a steep increase in computational power and VRAM demand for both LSTM and transformer encoding architectures when compared to 1D CNNs. As an illustration, a version of DeepMDC for the *K. pneumoniae - meropenem* pair was trained on a H100 GPU with 80GB VRAM. It was possible to attain a LSTM layer with 24 units only, having slightly inferior performance when compared to the DeepMDC version using the 1D CNN encoder.

A biology-informed extension of DeepMDC represents a promising direction for future work. Drawing inspiration from Physics-Informed Neural Networks (Cuomo et al., 2022), biological knowledge could be incorporated at multiple levels of the model. For example, known resistance gene families could be used to guide the attention mechanism, while genomic context, such as proximity to insertion sequences or plasmid backbones, could be used to modulate the relative importance of ORFs. In addition, phylogenetic relationships could be integrated into the loss function to discourage predictions that are inconsistent with known evolutionary structure.

Such approaches, recently explored in other domains such as cancer genomics (Wysocka et al., 2023), have the potential to improve performance in data-limited settings while enhancing the biological interpretability of model predictions.It is worth noting that classifying bacterial strains as resistant or susceptible to antibiotics is just one example of bacterial profiling that DeepMDC can perform. Other relevant applications include classifying strains by virulence level or biofilm-forming capacity. Future work will explore other bacterial profiling applications of DeepMDC and integrate additional layers of biological information, such as regulatory interactions, gene expression data, and epigenetic modifications. Incorporating multi-omics data may further enhance predictive accuracy and biological insight.

While models that use only genomic data as input effectively capture the presence of resistance markers, they may overestimate phenotypic resistance because they fail to account for transcriptional or post-transcriptional regulation. Another future direction for research is to augment the model to jointly analyze genomic and transcriptomic data. The architecture proposed in this work is well-suited to processing the augmented input set, thanks to the modern Hopfield network’s exponential storage capacity and the computational efficiency of 1D CNNs. One difficulty is the limited availability of both transcriptomic and genomic data for many strains. Also, the model relies on ORF prediction and translation, which may introduce biases or errors depending on annotation quality. And lastly, although attention weights provide interpretability, they do not necessarily imply causal relationships. Further experimental validation is required to confirm the biological relevance of identified features.In summary, the attention-based interpretability analysis provided valuable insights into the genomic signals that shape our deep learning model’s decisions for antimicrobial resistance prediction in *Klebsiella pneumoniae*. The model consistently prioritized biologically coherent features, including Tn*4401*-like transposases, *OmpK36* porins, and regulatory or phage-associated proteins. Nevertheless, attention-based models carry inherent interpretive limitations. Therefore, further studies are necessary to elucidate the role of these unannotated loci in resistance and virulence. Our findings reinforce the central role of Tn*4401* in the dissemination of carbapenemase genes and in modulating *OmpK36*-mediated permeability, a determinant of carbapenem resistance.

## Author Contributions

Nicolas da Matta Freire Araujo and Mateus Fernandes Santos developed DeepMDC and generated the results. Rafaela Correia Brum and Felipe Ramos Pinheiro performed data analysis. Renata Freire Alves Pereira, Bruno de Araújo Penna, Thiago Pavoni Gomes Chagas, Fábio Aguiar-Alves, Beatriz de Lima Alessio Müller, Audrien Alves Andrade de Souza, Alessandra Beatriz Santos Rondon Souza, Ágatha Ferreira de Souza, and Aline dos Santos Moreira performed sample collection, processing, and biologically oriented analysis of the results. Melise Chaves Silveira, Ana Paula D’Alincourt Carvalho-Assef, and Felicita Mabel Duré provided CABGen-based support and analysis. Marcelo Trindade dos Santos and Márcia da Silva Chagas contributed to the selection of evaluation metrics and assessments. Rafaela Correia Brum supported Amazon AWS executions. Fabricio Alves Barbosa da Silva and Adriano Maurício de Almeida Côrtes proposed the DeepMDC architecture. All authors participated in the discussion of the results. All authors contributed to the manuscript writing. All authors approved the final version of the manuscript.

## ACKNOWLEDGEMENTS

We acknowledge the INOVA FIOCRUZ program (VPPCB-002-FIO-20-2-23) for financial support. We thank CNPq for funding access to Amazon AWS (CNPq/AWS 421828/2022-6). We also acknowledge FAPERJ for financial support (MEDICINA DE PRECISÃO - E_20/2021). We would like to thank Bruna Jesuíno for her assistance in organizing and processing the training samples. We thank Rodolpho Albano for valuable discussions.

