## Supplemental tables for "Harnessing Interpretable Deep Learning to Predict Resistance in *Klebsiella pneumoniae*"

Table S1 - Validation and test set results for meropenem

| | Fold | ROC AUC | BACC | F1 score | MCC |
|---|---|---|---|---|---|
| Validation | 1 | 0,93 | 0,91 | 0,9 | 0,81 |
| Validation | 2 | 0,95 | 0,9 | 0,9 | 0,8 |
| Validation | 3 | 0,89 | 0,83 | 0,83 | 0,67 |
| Validation | 4 | 0,87 | 0,83 | 0,83 | 0,67 |
| Validation | 5 | 0,94 | 0,87 | 0,88 | 0,75 |
| Test | 1 | 0,82 | 0,81 | 0,78 | 0,63 |
| Test | 2 | 0,93 | 0,86 | 0,86 | 0,72 |
| Test | 3 | 0,75 | 0,74 | 0,72 | 0,49 |
| Test | 4 | 0,74 | 0,75 | 0,72 | 0,51 |
| Test | 5 | 0,8 | 0,74 | 0,75 | 0,47 |

Table S2 - Statistics for meropenem

| | | ROC AUC | BACC | F1 score | MCC |
|---|---|---|---|---|---|
| Average (Validation) | All Folds | 0,916 | 0,868 | 0,868 | 0,74 |
| Average (Test) | All Folds | 0,808 | 0,78 | 0,766 | 0,564 |
| Std Dev (Validation) | All Folds | 0,034 | 0,038 | 0,036 | 0,068 |
| Std Dev (Test) | All Folds | 0,076 | 0,053 | 0,058 | 0,107 |

Table S3 - Validation and test set results for cefepime

| | Fold | ROC AUC | BACC | F1 score | MCC |
|---|---|---|---|---|---|
| Validation | 1 | 0,79 | 0,71 | 0,74 | 0,42 |
| Validation | 2 | 0,71 | 0,66 | 0,71 | 0,32 |
| Validation | 3 | 0,78 | 0,72 | 0,76 | 0,43 |
| Validation | 4 | 0,81 | 0,73 | 0,76 | 0,45 |
| Validation | 5 | 0,79 | 0,73 | 0,76 | 0,46 |
| Test | 1 | 0,83 | 0,76 | 0,79 | 0,51 |
| Test | 2 | 0,8 | 0,74 | 0,76 | 0,47 |
| Test | 3 | 0,81 | 0,72 | 0,75 | 0,43 |
| Test | 4 | 0,78 | 0,69 | 0,72 | 0,37 |
| Test | 5 | 0,8 | 0,71 | 0,77 | 0,42 |

Table S4 - Statistics for cefepime

| | | ROC AUC | BACC | F1 score | MCC |
|---|---|---|---|---|---|
| Average (Validation) | All Folds | 0,776 | 0,71 | 0,746 | 0,416 |
| Average (Test) | All Folds | 0,804 | 0,724 | 0,758 | 0,44 |
| Std Dev (Validation) | All Folds | 0,038 | 0,029 | 0,022 | 0,056 |
| Std Dev (Test) | All Folds | 0,018 | 0,027 | 0,026 | 0,053 |

Table S5 - Validation and test set results for ceftazidime

| | Fold | ROC AUC | BACC | F1 score | MCC |
|---|---|---|---|---|---|
| Validation | 1 | 0,94 | 0,87 | 0,92 | 0,66 |
| Validation | 2 | 0,93 | 0,88 | 0,93 | 0,67 |
| Validation | 3 | 0,92 | 0,87 | 0,93 | 0,67 |
| Validation | 4 | 0,92 | 0,87 | 0,9 | 0,63 |
| Validation | 5 | 0,96 | 0,88 | 0,92 | 0,67 |
| Test | 1 | 0,88 | 0,88 | 0,94 | 0,72 |
| Test | 2 | 0,9 | 0,83 | 0,92 | 0,61 |
| Test | 3 | 0,91 | 0,85 | 0,93 | 0,67 |
| Test | 4 | 0,92 | 0,85 | 0,9 | 0,6 |
| Test | 5 | 0,92 | 0,87 | 0,93 | 0,68 |

Table S6 - Statistics for ceftazidime

| | | ROC AUC | BACC | F1 score | MCC |
|---|---|---|---|---|---|
| Average (Validation) | All Folds | 0,934 | 0,874 | 0,92 | 0,66 |
| Average (Test) | All Folds | 0,906 | 0,856 | 0,924 | 0,656 |
| Std Dev (Validation) | All Folds | 0,017 | 0,005 | 0,012 | 0,017 |
| Std Dev (Test) | All Folds | 0,017 | 0,019 | 0,015 | 0,050 |

Table S7 - Validation and test set results for gentamicin

| | Fold | ROC AUC | BACC | F1 score | MCC |
|---|---|---|---|---|---|
| Validation | 1 | 0,94 | 0,89 | 0,89 | 0,78 |
| Validation | 2 | 0,9 | 0,85 | 0,84 | 0,71 |
| Validation | 3 | 0,89 | 0,83 | 0,82 | 0,66 |
| Validation | 4 | 0,96 | 0,89 | 0,89 | 0,79 |
| Validation | 5 | 0,9 | 0,82 | 0,82 | 0,64 |
| Test | 1 | 0,89 | 0,81 | 0,81 | 0,62 |
| Test | 2 | 0,91 | 0,75 | 0,77 | 0,51 |
| Test | 3 | 0,74 | 0,71 | 0,73 | 0,42 |
| Test | 4 | 0,86 | 0,75 | 0,75 | 0,51 |
| Test | 5 | 0,89 | 0,8 | 0,8 | 0,59 |

Table S8 - Statistics for gentamicin

| | | ROC AUC | BACC | F1 score | MCC |
|---|---|---|---|---|---|
| Average (Validation) | All Folds | 0,918 | 0,856 | 0,852 | 0,716 |
| Average (Test) | All Folds | 0,858 | 0,764 | 0,772 | 0,53 |
| Std Dev (Validation) | All Folds | 0,030 | 0,033 | 0,036 | 0,068 |
| Std Dev (Test) | All Folds | 0,068 | 0,041 | 0,033 | 0,078 |
